# Sex-dependent differences in pain and sleep in a porcine model of Neurofibromatosis type 1

**DOI:** 10.1101/495358

**Authors:** Rajesh Khanna, Aubin Moutal, Katherine A. White, Aude Chefdeville, Pedro Negrao de Assis, Song Cai, Vicki J. Swier, Shreya S. Bellampalli, Marissa D. Giunta, Benjamin W. Darbro, Dawn E. Quelle, Jessica C. Sieren, Margaret R. Wallace, Christopher S. Rogers, David K. Meyerholz, Jill M. Weimer

## Abstract

Neurofibromatosis Type 1 (NF1) is an autosomal dominant genetic disorder resulting from germline mutations in the *NF1* gene, which encodes neurofibromin. Patients experience a variety of symptoms, but pain in the context of NF1 remains largely underrecognized. Here, we characterize nociceptive signaling and pain behaviors in a miniswine harboring a disruptive *NF1* mutation (exon 42 deletion). We explore these phenotypes in relationship to collapsin response mediator protein 2 (CRMP2), a known interactor of neurofibromin. Mechanistically, we found two previously unknown phosphorylated residues of CRMP2 in *NF1^+/ex42del^* pig dorsal root ganglia (DRGs) and replicated increased voltage-gated calcium channel currents in *NF1^+/ex42del^* pig DRGs previously described in rodent models of NF1. We present the first characterization of pain-related behaviors in a pig model of NF1, identifying unchanged agitation scores, lower tactile thresholds (allodynia), and decreased response latencies to thermal laser stimulation (hyperalgesia) in the *NF1* mutant animals; *NF1^+/ex42del^* pigs demonstrated sexually dimorphic behaviors. *NF1^+/ex42del^* pigs showed reduced sleep quality and increased resting, two health-related quality of life symptoms found to be comorbid in people with NF1 pain. Finally, we show decreased depolarization-evoked calcium influx in both wildtype and *NF1^+/ex42del^* pig DRGs treated with CRMP2 phosphorylation inhibitor (5)-lacosamide. Our data supports use of *NF1^+/ex42del^* pigs as an ideal model for studying NF1-associated pain and are a better model for understanding the pathophysiology of NF1 compared to rodents. Moreover, our findings demonstrate that interfering with CRMP2 phosphorylation might be a promising therapeutic strategy for NF1-related pain management.

## 1. Introduction

Neurofibromatosis type 1 (NF1) is a human autosomal dominant genetic disorder resulting from germline mutations in the *NF1* gene [8; 20; 50; 51]. The *NF1* gene on chromosome 17q11.2 encodes the 320kDa cytoplasmic protein Neurofibromin. Neurofibromin is ubiquitously expressed during development [12] and highly expressed only in nervous system tissues during adulthood [13]. Neurofibromin acts as a negative regulator of the protooncogene Ras activity through its RAS-GTPase activating domain [2; 58]. Therefore, impaired neurofibromin function increases Ras activation and stimulates downstream RAF/MEK and AKT/mTOR signaling pathways [11]. Neurofibromin also acts as an adenyl cyclase activity regulator [48], and may have other functions not yet characterized.

People with NF1 can experience a variety of symptoms including, pigmentary abnormalities (café-au-lait macules, axillary/inguinal freckling) [23], skeletal deformities, cardiovascular abnormalities, and cognitive deficits (attention and learning impairments) [24]. Individuals with NF1 are prone to develop benign, and occasionally malignant, tumors of the central and peripheral nervous system. About 15% of children with NF1 develop a glioma of the optic pathway [29] and adults bear a 50-100-fold increased risk of developing a malignant glioma compared to the general population [21]. NF1 patients are also at high risk for the development of malignant peripheral nerve sheath tumors (MPNSTs), which are the leading cause of death in this patient population [44]. Malignancies outside of the nervous system, such as leukemias, are also seen [44]. Finally, patients with this disease report experiencing abnormal levels of pain [3] but chronic pain in the context of NF1 is largely underrecognized and understudied.

In the 1990s, mouse strains harboring mutations in *Nf1* were created to model NF1, as there are no naturally-occurring animal models of NF1. Heterozygous *Nf1^+/-^* mice presented with moderate learning and memory impairments [45] but did not develop hallmarks of the disease such as spontaneous tumors nor exhibit pigmentation defects [26]; the latter required engineered loss of the wildtype allele in specific cell types. CNS tumors were only observed following co-inactivation of tumor suppressor genes, such as *p53* or Phosphatase and tensin homolog *(Pten)*, in neuroglial precursors [27; 60]. Thus, transgenic mouse models are valuable for studying select clinical manifestations of NF1 but fail to mimic the spectrum of disease features of patients.

To design a better NF1 animal model, a miniswine harboring a disruptive *NF1* mutation (exon 42 deletion) was engineered [55]. This miniswine spontaneously exhibited café-au-lait macules, learning impairment and neurofibromas, thus recapitulating classic features of NF1, many of which are not displayed in conventional heterozygotic rodent models [55]. However, pain perception and signaling in NF1 have not been investigated. We have previously shown that collapsin response mediator protein 2 (CRMP2) is necessary for NF1-related pain [34; 38]. Here, we have developed novel tools to monitor behaviors related to pain in NF1-mutant miniswine. Moreover, we characterize the expression of CRMP2 and study the regulation of nociceptive calcium channel function by CRMP2 in this novel pig model. We suggest that pain modeling in large animals, such as pigs, offers considerable translational potential for driving therapeutic discoveries.

## 2. Methods

### 2.1. Immunofluorescence

Porcine DRGs were harvested and fixed in 4% PFA for 24 hours, cryoprotected in 30% sucrose (m/v in PBS) for 48 hours and frozen at −70°C in 2-methylbutane chilled with dry ice. Samples were cut into 20μm-thick sections on a cryostat. Sections were subsequently blocked in 3% BSA, 0.1% Triton X-100 in PBS for 1 hour at room temperature and incubated with primary antibody diluted in blocking solution overnight at +4°C. After washes in PBS, slides were incubated with secondary antibody diluted in blocking solution for 2 hours at room temperature, washed and counterstained with DAPI. For the negative control, antibody was omitted in the protocol. Images were acquired on an Axio Observer.Z1 (Zeiss), using objectives EC Plan-Neofluar 10x (NA 0.30) and controlled by the Zen software (Zeiss).

### 2.2. Culturing of miniswine DRG neurons

DRG neurons were isolated from 7-9-month-old Yucatan miniswine using methods described previously [55]. At necropsy, a bilateral laminectomy was made and DRGs were collected then digested in 3 ml neurobasal media (cat# 21103049, Thermofisher Scientific) containing collagenase Type II (Cat# LS004205, Thermofisher, 12 mg/ml) and dispase II (Cat# 17-105-041, Thermofisher, 20 mg/ml) and incubated for 45-75 minutes at 37°C under gentile agitation. Dissociated DRG neurons (~1.5 × 10^6^) were then gently centrifuged to collect cells and washed with and grown in Neurobasal media containing 1% penicillin/streptomycin sulfate from 10,000 μg/ml stock, 30 ng.ml^-1^ nerve growth factor (Thermofisher), 10% FBS (Hyclone) and 2% B-27 (Cat# 17-504-044, Thermofisher).

### 2.3. Calcium imaging in acutely dissociated miniswine DRG neurons

Dorsal root ganglion neurons were loaded for 30 minutes at 37°C with 3 μM Fura-2AM (Cat# F1221, Thermo Fisher, stock solution prepared at 1mM in DMSO, 0.02% pluronic acid, Cat#P-3000MP, Life Technologies) to follow changes in intracellular calcium([Ca^2+^]_c_) in a standard bath solution containing 139 mM NaCl, 3 mM KCl, 0.8 mM MgCl_2_, 1.8 mM CaCl_2_, 10 mM Na HEPES, pH 7.4, 5 mM glucose exactly as previously described [5] Fluorescence imaging was performed with an inverted microscope, NikonEclipseT*i*-U (Nikon Instruments Inc., Melville, NY), using objective Nikon Fluor 4X and a Photometrics cooled CCD camera Cool SNAP ES^2^ (Roper Scientific, Tucson, AZ) controlled by Nis Elements software (version 4.20, Nikon Instruments). The excitation light was delivered by a Lambda-LS system (Sutter Instruments, Novato, CA). The excitation filters (340 ± 5 and 380 ± 7) were controlled by a Lambda 10 to 2 optical filter change (Sutter Instruments). Fluorescence was recorded through a 505-nm dichroic mirror at 535 ± 25 nm. To minimize photobleaching and phototoxicity, the images were taken every ~10 seconds during the time-course of the experiment using the minimal exposure time that provided acceptable image quality. The changes in [Ca^2+^]_c_ were monitored by following a ratio of F_340_/F_380_, calculated after subtracting the background from both channels.

### 2.4. Whole-cell patch recordings of Ca^2+^ currents in acutely dissociated miniswine DRG neurons

Recordings were obtained from acutely dissociated DRG neurons as described previously [25; 37]. To isolate calcium currents, Na^+^ and K^+^ currents were blocked with 500 nM tetrodotoxin (TTX; Alomone Laboratories) and 30 mM tetraethylammonium chloride (TEA-Cl; Sigma). Extracellular recording solution (at ~310 mOsm) consisted of the following (in mM): 110 *N*-methyl-D-glucamine (NMDG), 10 BaCl_2_, 30 TEA-Cl, 10 HEPES, 10 glucose, pH at 7.4, 0.001 TTX, 0.01 nifedipine. The intracellular recording solution (at ~310 mOsm) consisted of the following (in mM): 150 CsCl_2_, 10 HEPES, 5 Mg-ATP, 5 BAPTA, pH at 7.4. Activation of I_Ca_ was measured by using a holding voltage of – 90 mV with voltage steps 200 milliseconds in duration applied at 5-second intervals in + 10 mV increments from −70 to +60 mV. Current density was calculated as peak I_Ca_/cell capacitance. Steady-state inactivation of I_Ca_ was determined by applying an 800-millisecond conditioning prepulse (−100 to −20mV in +10mV increments) after which the voltage was stepped to −20 mV for 200 milliseconds; a 15-second interval separated each acquisition. Whole-cell voltage clamp recordings were performed at room temperature using an EPC 10 Amplifier-HEKA as previously described [15]. The internal solution for voltage clamp sodium current recordings contained (in mM): 140 CsF, 1.1 CsEGTA, 10 NaCl, and 15 HEPES (pH 7.3, 290-310 mOsm/L) and external solution contained (in mM): 140 NaCl, 3 KCl, 30 tetraethylammonium chloride, 1 CaCl_2_, 0.5 CdCl_2_, 1 MgCl_2_, 10 D-glucose, and 10 HEPES (pH 7.3, 310-315 mosM/L).

Fire-polished recording pipettes, 2 to 5 MΩ resistance were used for all recordings. Whole-cell recordings were obtained with a HEKA EPC-10 USB (HEKA Instruments Inc., Bellmore, NY); data were acquired with a Patchmaster (HEKA) and analyzed with a Fitmaster (HEKA). Capacitive artifacts were fully compensated, and series resistance was compensated by ~70%. Recordings made from cells with greater than a 5mV shift in series resistance compensation error were excluded from analysis. All experiments were performed at room temperature (~23°C).

The Boltzmann relation was used to determine the voltage dependence for activation of ICa and wherein the conductance-voltage curve was fit by the equation G/G_max_ = 1/ [1 + exp (V_0.5_ – V_m_)/k], where G is the conductance G=I/(V_m_-E_Ca_), G_max_ is the maximal conductance obtained from the Boltzmann fit under control conditions, V_0.5_ is the voltage for half-maximal activation, V_m_ is the membrane potential, and k is a slope factor. E_Ca_ is the reversal potential for I_Ca_; E_Na_ is the reversal potential for I_Na_ and was determined for each individual neuron. The values of I_Ca_ and around the reversal potential were fit with a linear regression line to establish the voltage at which the current was zero. The Boltzmann parameters were determined for each individual neuron and then used to calculate the mean ± S.E.M.

### 2.5. Assessment of Spontaneous Pain Behavior Using a Composite Behavior Scale and Home Pen Monitoring

The solitary performance and social behavior of each animal was scored during a 10-minute observation period as described previously [7]. Seven behavioral parameters were observed and recorded (3 for solitary performance and 4 for social behavior). The animals were observed in their home pen animals and are monitored bi-monthly for indicators of pain, anxiety, hyperactivity, and tumor growth. These bi-monthly observations (by the caretaker who handled them from the first acclimation day) included: monitoring of café au lait spots, tumor growth, blindness, loss of vision, body weight, body condition, seizures, limb paralysis, lethargy, feeding behavior, ability to bear body weight, vocalization, restlessness, agitation, aggression, and isolation. Score driving parameters related to observing the animals’ standing posture, leg guarding, leg shaking, as well as their vocalization and social behavior (isolation and aggressiveness). Each parameter was graded from 0 to 2, depending on the observed behavior. The sum of all points from the 7 parameters was considered as the final score. A higher score indicates that the animal expressed more spontaneous pain behavior. The maximum possible score was 11 points. Spontaneous expression scores were observed monthly or biweekly as permissible (Adapted from [7]). In addition, the pigs were subjected to whole body MRI to identify any internal tumors.

Additionally, each animal was tracked for 6 days using a FitBark2 (www.fitbark.com) device, attached to their neck using a common dog collar at 10-12 months of age. Readouts included: sleep quality, time spent being active, playing, and resting, distance travelled, and weight. This allowed us to monitor the animals’ activities without the influence of human interaction. At this time point, female wildtype animals were not available for testing, so female NF1 mutant animals were compared to male wildtype animals.

### 2.6. Assessment of Tactile Allodynia

In order to gauge an animal’s level of pain/sensitivity, we monitored the animal’s response to von Frey sensory probes (Stoelting 58011; probe range 0.008-300 grams) when stimulated on the skin of the anthelix (a part of the visible ear; the pinna. The antihelix is a curved prominence of cartilage parallel with and in front of the helix on the pinna) just inside the ear. Animals were monitored for response to sensory probes over five consecutive trials stating at 1 gram of target force, with a positive response deemed as the animal responding at least three out of five trials (ear flick, facial muscle twitch, or similar response). If the animal responded to the 1-gram target force, lower pressure probes were used to stimulate the anthelix of the ear until at five trials each, until the lowest target force that garnished a response was determined. If the animal did not respond to the 1-gram target force, higher pressure probes were used to stimulate the anthelix of the ear until at five trials each, until the highest target force that garnished a response was determined. The lowest target force detected by each animal was converted on a log2 scale (as the target probes are non-linear and animals perceive stimuli on a logarithmic scale (Weber’s Law)) [33].

### 2.7. Thermal Laser Stimulation

Withdrawal reflex via thermal laser stimulation was modified based on Herskin et al. [22]. Animals were secured in a sling and a 2W S3 Arctic laser (Wicked Laser; http://www.wickedlasers.com/arctic) was used to stimulate three distinct areas of the back of the outer ear for three, 20 second trials at a distance of 100cm (see Figure 4A). With each distinct area, the latency to respond to the laser stimulation was recorded, and the latency to respond to each trial was averaged into one value per animal. The original Herskin et al publication [22] stimulated the shoulder and leg, though we found our animals would not consistently respond to stimulation in those areas (data not shown).

### 2.8. Additional pain assessment protocols

Bright light, pigeon feather, and air puff tests were employed in an attempt to measure stimuli response in swine subjects secured in a sling.

#### 2.8.1. Bright Light Exposure

An LED headlamp was affixed to the experimenter’s head (Amazon B00H26CGTI) in a dark room, and the animal was exposed to a bright, flashing strobe light for five 15 second trials. Avoidance response to the light was recorded.

#### 2.8.2. Pigeon Feather Stimulation

A single, natural, 6-8-inch-long pigeon feather (Amazon B071JF1JFC) was gently brushed across various areas of the animal’s face, such as the snout and undereye area, where skin was anticipated to be thinnest. The response out of five trials was recorded.

#### 2.8.3. Air Puff Stimulation

A 10cc syringe was placed 8-10 cm from the side of the animal’s eye and a puff of air was directed at the eyelids. The response out of five trials was recorded.

### 2.9. Study approval

All animals were maintained at Exemplar Genetics under an approved IACUC protocol (protocol no. MRP2016-009). DRG samples were obtained from individuals targeted for necropsy and sample were shared with multiple other labs at the University of Iowa in accordance with the IACUC approved protocol (Office of Animal Resources, University of Iowa; protocol no. 7061269).

### 2.10. Experimental Design and Statistical Measurement

All animals were randomly assigned to distinct experimental groups by an off-site experimenter. All technicians performing the behavior experiments were blinded to genotype, with all data analyzed by a blinded, off-site experimenter. Statistical tests are noted in the figure legends and include an unpaired student’s t-test (pooled sexes) and an ordinary two-way ANOVA with a Tukey post-hoc correction (split sexes). Graphs are represented as scatter plots with mean ± SEM.

## 3. Results

### 3.1. NF1 mutant miniswine generation and characterization of ion channel remodeling

We recently reported the generation of *NF1^+/ex42del^* Yucatan miniature swine via recombinant adeno-associated virus-mediated gene targeting and somatic cell nuclear transfer methods [55]. As in patients with NF1, all affected swine were heterozygous. *NF1^+/ex42del^* animals presented with spontaneous café au lait macules and neurofibromas, two classic disease phenotypes that are absent in heterozygous *Nf1* rodent models [55]. The *NF1^+/ex42del^* miniswine also presented with initial learning delay in the T-maze task, an initial hesitance to interact with a novel object, and overall observations of hyperactivity and anxiety [55]. Since previous reports have demonstrated an increase in voltage gated N-type Ca^2+^ (CaV2.2) channel current densities in neurons from *Nf1^+/-^* mice, placing CaV2.2 as the major contributor to whole cell Ca^2+^ currents in NF1, with no significant differences in L-, P/Q-, T- and R-type currents [14; 40], we used whole-cell voltage-clamp electrophysiology to assess the activity of these channels in DRGs from wildtype and *NF1^+/ex42del^* pigs. Representative recordings from a sensory neuron from wildtype and *NF1^+/ex42del^* pigs are illustrated in Figure 1A.

**Figure 1.**
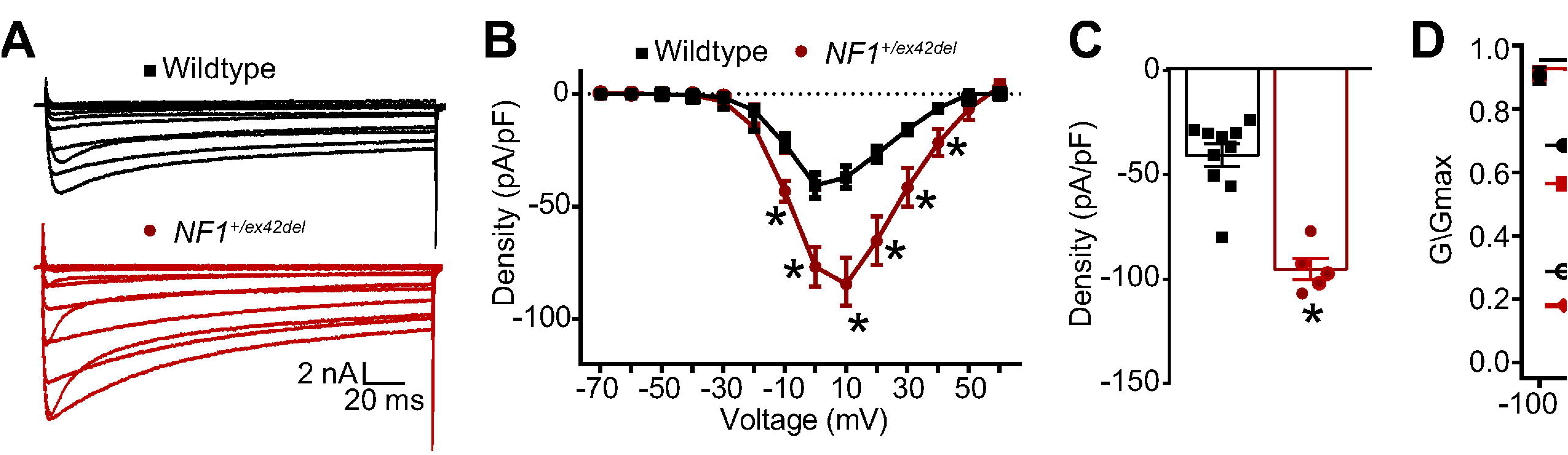
Calcium currents are increased in *NF1^+/ex42del^* pig DRGs. (**A**) representative calcium traces from wildtype or *NF1^+/ex42del^* pig DRGs. (**B**) Summary current-voltage curve of peak current density normalized to the cell capacitance (pA/pF) for each voltage step. (**C**) Bar graph with scatter plot of peak current density at 10 mV for the indicated conditions. (D) Activation and inactivation curves of normalized calcium currents (G/Gmax) for the indicated genotype. *p<0.05, One-way ANOVA with Tukey post-hoc test, n= 6-10 cells per condition. Unless otherwise staed, all data are presented as means ± SEM.

To account for the variations in cell size, the calcium currents obtained from 10 wildtype and 5 *NF1^+/ex42del^* sensory neurons were normalized to cell capacitance and shown as the current density-voltage relation summarized in Figure 1B. The cell capacitance for the isolated neurons was not different between the two genotypes (data not shown). There was an ~2.3-fold significant increase in the average peak current densities between wildtype compared to *NF1^+/ex42del^* neurons (−40.6 ± 5.4 vs. −94.3 ± 4.5 pA/pF, respectively, for steps to +10 mV, p<0.05, t-test) (Figure 1B, C). When the current values were transformed to conductance (G), the conductance-voltage relation was fit with the Boltzmann relation, and the conductance for each neuron was then normalized to the maximal value of G (Gmax) obtained from the fit. The G/Gmax-voltage relation is summarized in Figure 1D and indicates that the voltage-dependence for activation of Gmax was nearly identical between the two genotypes: the Boltzmann fitting parameters, voltage for half-maximal activation (V_0.5_) and slope *(k)* were −26.3 ± 1.3 and 8.8 ± 1.1 for wildtype neurons, respectively, and −24.9 ± 1.5 and 9.1 ± 1.5 for *NF1^+/ex42del^* neurons, respectively. Similarly, the voltage-dependence for inactivation of Gmax was nearly identical between the two genotypes: V_0.5_ and *k* were −6.9 ± 1.0 and 6.0 ± 0.8 for wildtype neurons, respectively, and −9.2 ± 1.6 and 4.5 ± 1.1 for *NF1^+/ex42del^* neurons, respectively. These results confirmed that calcium currents are increased in *NF1^+/ex42del^* pig DRGs compared to wildtype swine DRGs.

### 3.2. Changes in CRMP2 phosphorylation in NF1^+/ex42del^ pig DRGs underpin the changes in calcium channel remodeling

To determine the mechanism of the increased calcium currents in *NF1^+/ex42del^* neurons, we explored the underlying signaling. In neurons, neurofibromin interacts with CRMP2 to repress its phosphorylation by cyclin dependent kinase 5 (Cdk5) on a serine residue at position 522 of CRMP2 – a site found to be important for hyperalgesia in the context of NF1 [34; 38; 43]. We have also reported that CRISPR/Cas9 mediated excision of the C-terminal domain of neurofibromin abolished the interaction with CRMP2, thus leading to increased CRMP2 phosphorylation in rodent DRG and spinal cord [41]. We asked if this effect on CRMP2 phosphorylation is recapitulated in our new *NF1* mutant miniswine model. We used immunofluorescence staining to assess CRMP2 expression level and phosphorylation in the DRGs from wildtype and *NF1^+/ex42del^* pigs. DRG slices were co-stained for either anti-CRMP2 or anti-CRMP2 p522 and anti-βIII-tubulin (to identify neurons) antibodies. Specific staining was observed for CRMP2 and p522 CRMP2 in pig DRG neurons, thus showing that our antibodies are amenable to detect CRMP2 and its phosphorylated form in pig (Figure 2A).

**Figure 2.**
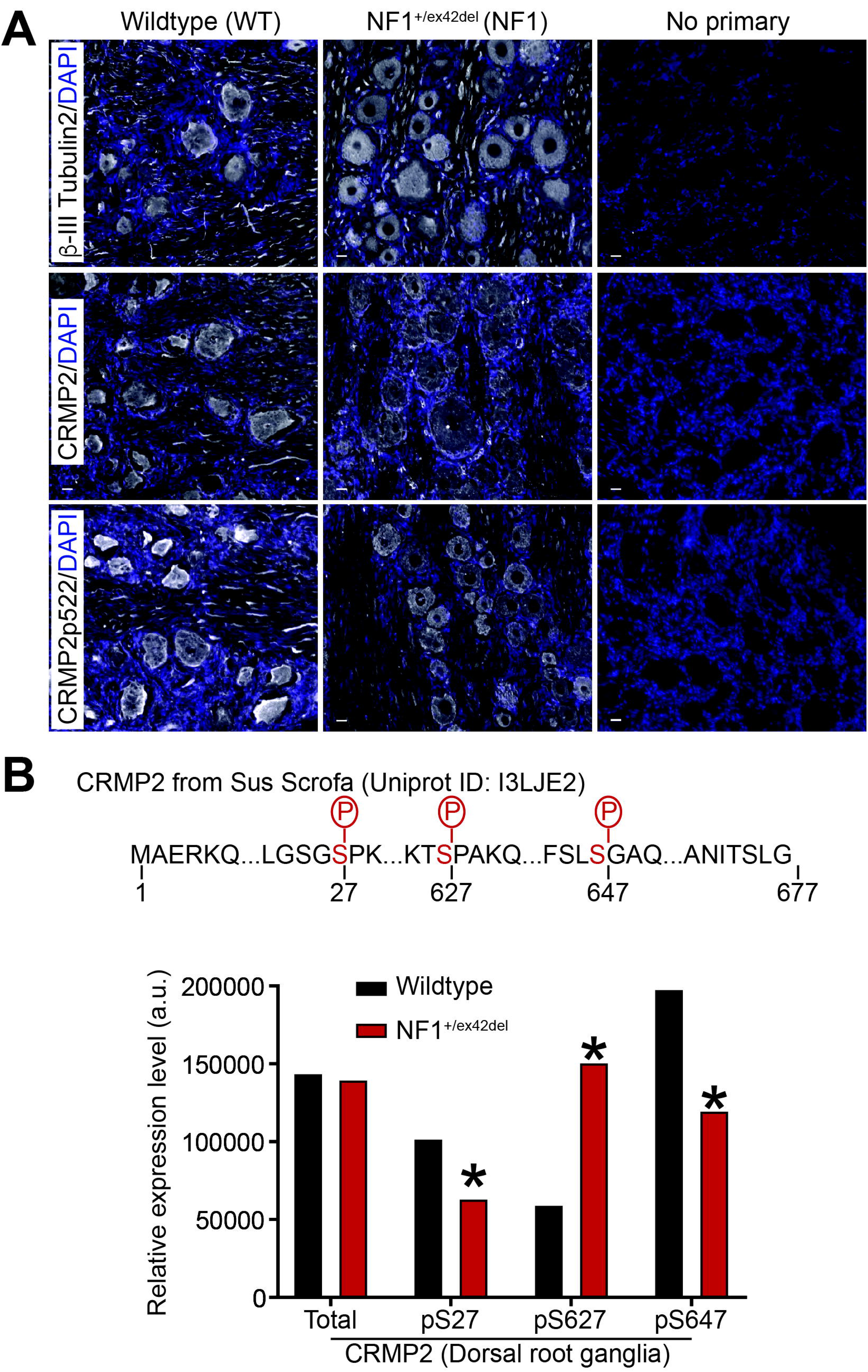
Phosphoproteomic analysis of CRMP2 phosphorylation in *NF1^+/ex42del^* pig DRGs. (**A**) immunostainings on PFA-fixed pig DRG slices from WT or NF1 *^+/ex42del^* pigs. Nuclei were counterstained with DAPI. For the negative control, primary antibody was omitted. Antibodies against CRMP2 phosphorylation sites p509/p514 (phosphorylation by GSK3β) or p555 (phosphorylation by RHOK) did not yield any specific signal (data not shown). Scale bars: 20μm. (**B**) Wildtype or *NF1^+/ex42del^* pig DRGs whole lysate was analyzed by mass spectrometry for CRMP2 expression and phosphorylation level. The peptide ions m/z was analyzed based on CRMP2 protein sequence from pigs *(sus scrofa*) in Uniprot (I3LJE2). Partial CRMP2 protein sequence is shown and the identified phosphorylation sites highlighted in red. Mass spectrometry returned a quantitative value of peptide abundance compared to the sample load, presented in the bar graph. CRMP2 total expression was unchanged in *NF1^+/ex42del^* compared to wildtype pig DRGs. Expression level of the phosphorylation sites pS27, pS627 and pS647 was altered in *NF1^+/ex42del^* pig DRGs. *p<0.05, Mann-Whitney test (n=6 animals per genotype). Error bars are smaller than the symbols.

Next, we used phosphoproteomics to assess CRMP2 phosphorylation in pig DRGs (Figure 2B). This is an unbiased approach that enables detection of all CRMP2 phosphorylation events independently of available antibodies, many of which have not been validated in pigs (data not shown). Using mass spectrometry, we analyzed whole DRG lysates from wildtype and *NF1^+/ex42del^* pigs (n=6 each). While CRMP2 expression levels were not changed by *NF1* mutation (Figure 2B), these analyses uncovered several novel observations about the porcine protein. First, we detected a long form of CRMP2 in pig DRGs. This long form results from a splicing event on the mRNA that adds nearly 100 amino acids onto the N-terminus of the protein. Prior studies showed that this alternate CRMP2 form exists in humans [47], but it is not present in rodents. The extensive genome and transcriptome sequencing performed in the last 20 years allows us to mine experimental evidence of transcripts for any gene. In the Ensembl and Nucleotide databases, alternative transcripts leading to a long form of CRMP2 are referenced for humans with published evidence at the protein level [47].

Second, we identified 3 phosphorylation sites on CRMP2 (Serine 27, Serine 627, and Serine 647) whose modification was altered by *NF1* status (Figure 2B). In DRGs of *NF1^+/ex42del^* pigs, S27 and S647 phosphorylation levels were decreased while the S627 phosphorylation level was increased (Figure 2B). The kinases responsible for phosphorylation at residues S27 and S647 are unknown. We used GPS 3.0 to predict the kinases (selected for a high threshold, false discovery rate <10%) that might mediate the phosphorylation of CRMP2 at these novel sites. Several potential kinases are implicated for S27 phosphorylation including those from the Dual specificity Tyrosine Regulated Kinase (DYRK), Cyclin Dependent Kinase (CDK) and Tau-Tubulin Kinase (TTBK) families but also by MAPK12 (Mitogen-activated protein kinase 12), and KIS, a kinase associated with microtubule regulators. The S647 phosphorylation site had a completely different predicted kinase landscape with candidate kinases including NIMA (never in mitosis gene a)-related kinase 2 (NEK2), LIM domain kinase 1 (LIMK1) and type 1 serine/threonine kinase receptors including activin-like receptors 1-7 (ALK1-7) were possible candidates for the phosphorylation. Serine 627 is analogous to the Cdk5 regulated S522 phosphorylation site that is present in both rodents and humans. These analyses identified two novel phosphorylation sites in CRMP2 and demonstrated that DRGS from *NF1^+/ex42del^* pigs have altered CRMP2 phosphorylation.

### 3.3. NF1^+/ex42del^ pigs exhibit reflexive allodynia and hyperalgesia as well as spontaneous pain behaviors in a sex- and time-dependent manner

People with NF1 report idiopathic pain [10] and we found that *Nf1* gene editing could directly result in allodynia in rats [41]. We asked if this was also true in our *NF1* mutant miniswine model. Since pain behavior testing in pigs has not been extensively attempted or studied, we first tried to replicate some of the classical rodent testing approaches for pain in the pigs. To this end, we employed a variety of techniques, including latency to move away from a bright, flashing light; time to react to a puff of air directed at the eye; tactile sensitivity test using a pigeon feather stimulus [7]; and reaction to von Frey filaments just inside the ear. The flashing light, air puff, and pigeon feather experiments were unsuccessful, as all animals regardless of genotype reacted extremely quickly and similarly to the flashing light and air puff tests, and animal did not respond to the pigeon feather experiment (data not shown). In contrast, we were able to successfully measure the reaction to the von Frey filaments. We observed a decrease in withdrawal threshold to von Frey filaments in tumor-burdened male *NF1^+/ex42del^* pigs at 7 and 8 months of age (Figure 3A) and in female *NF1^+/ex42del^* pigs at 7 and 8 months of age compared to age-and gender-matched wildtype pigs (Figure 3B). Looking at all pigs regardless of sex and tumor burden, combined data shows that *NF1^+/ex42del^* pigs have significant allodynia at 8 months (Figure 3C). Taken together, this dataset shows that *NF1^+/ex42del^* pigs have features of hypersensitivity and pain, as it has been reported by NF1 patients [49] and in rodent models of NF1 [39; 42; 56]. Importantly, allodynia in NF1 mutant pigs is sex and tumor burden dependent and decreases as the animals age.

**Figure 3.**
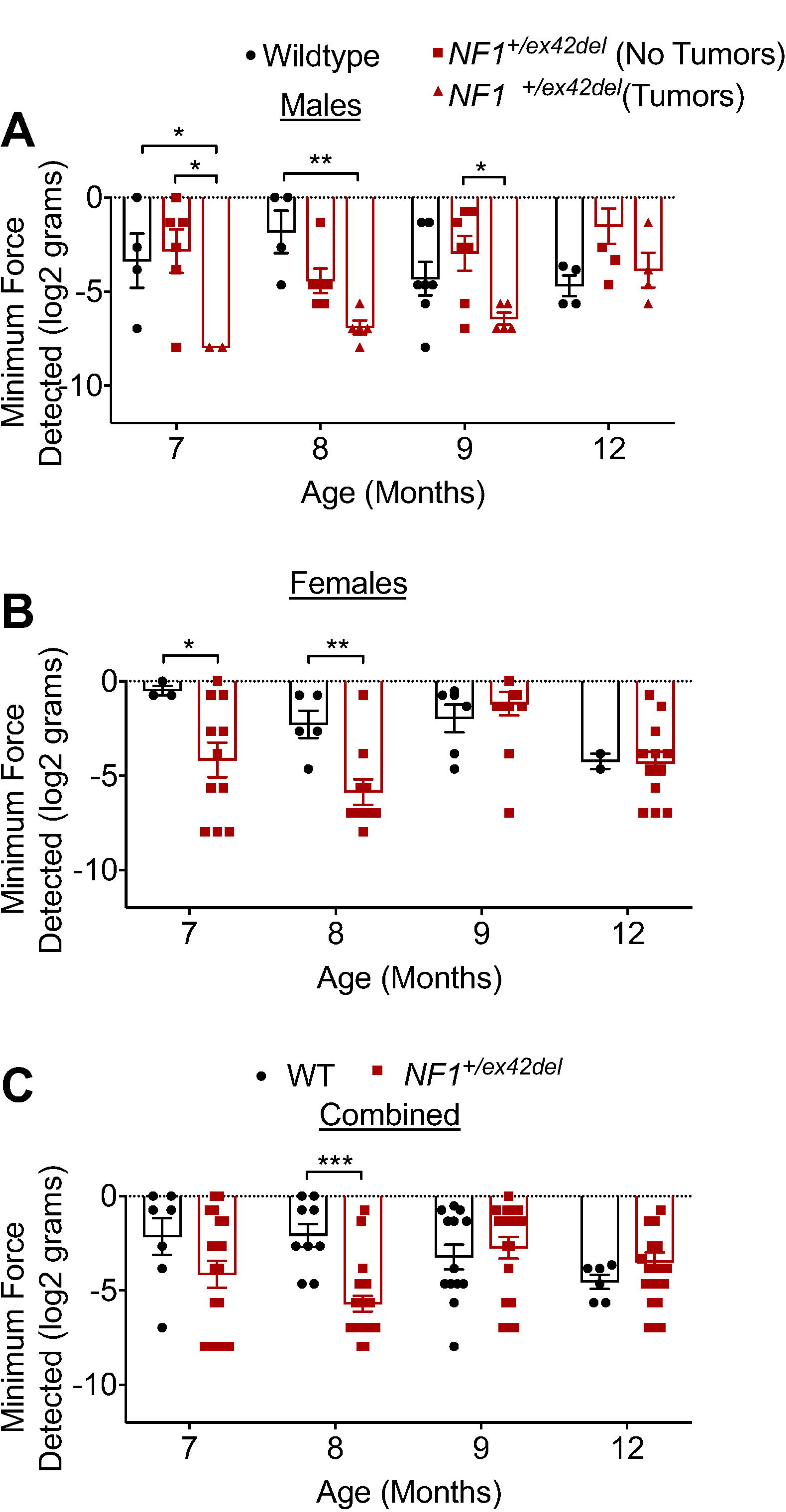
*NF1^+/ex42del^* pigs exhibit mechanical allodynia. Bar graph with scatter plot of inner ear tactile threshold values for male (A) or female (B) pigs (data is combined in (C)) from the indicated genotype. Tactile thresholds were decreased in both tumor-burdened male and female *NF1^+/ex42del^* pig (*p<0.05, **p<0.01, ***p<0.001. Two-way ANOVA with Tukey post-hoc.).

Next, we used a laser-based method to measure thermal nociception [22] in group-housed wildtype and *NF1^+/ex42del^* pigs. Withdrawal reflex of the hindlegs and shoulders was recorded in response to a thermal stimulation with a handheld laser with 1-2 watts power output. This response did not differ between the genotypes (data not shown). The nociceptive response of the pigs, assessed by examining their reactions to a laser beam applied to the outer part of the ear (Figure 4 A), resulted in reflexive behavioral responses in all animals tested. Male wildtype and *NF1^+/ex42del^* pigs (with and without neurofibromas) (Table 1) at 15-20 months of age did not show any differences in their latencies to respond to the laser shone on the ear (Figure 4B). Neurofibromas were only observed in males, on their upper body (Table 1) where the animals tended to ram into each other or rub aggressively against each other or their pens. In contrast, female *NF1^+/ex42del^* pigs had significantly faster response times to the laser compared to age-matched wildtype female pigs (Figure 4C), further supporting sexual dimorphism in NF1 swine model pain perception.

**Figure 4.**
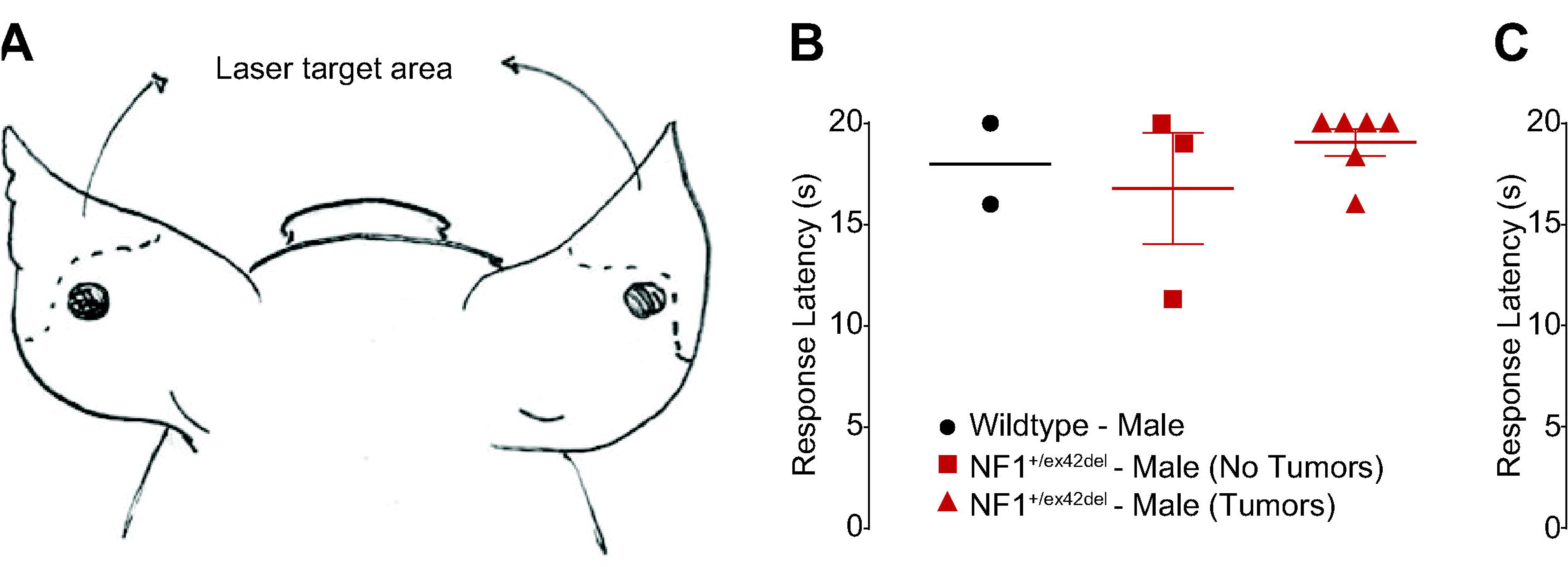
*NF1^+/ex42del^* pigs show sexually dimorphic behaviors in response to thermal laser stimulation. (**A**) Schematic of site of thermal laser stimulation in pigs. (**B**) Male *NF1^+/ex42del^* pigs exhibit non-significant differences in response latency to thermal laser stimulation in comparison to wildtype pigs regardless of tumor presence at 15 months of age. (**C**) Female *NF1^+/ex42del^* pigs exhibit significantly reduced response latencies to thermal laser stimulation in comparison to wildtype pigs at 15-20 months of age (*p<0.05, Mann Whitney for each age).

We also assessed spontaneous pain behaviors using a composite behavior scale reported by Meilin and colleagues [7]. In general, the parameters relate to observing the animals’ standing posture, leg guarding, leg shaking, as well as their vocalization and social behavior (isolation and aggressiveness). In this manner, the solitary performance and social behavior of each animal were scored during a 10-minute observation period. There were no differences between genotypes in weight bearing, appearance, vocalization, restlessness, aggression, and isolation scores, as all animals scored 0 in these parameters (data not shown). Starting at 6 months of age through 10 months of age, animals did score in markers of agitation, however there were no differences in the agitation scores of *NF1^+/ex42del^* pigs, either males (Figure 5A) or females (Figure 5B), compared to their age- and gender-matched wildtype counterparts. It is unclear if agitation is related to pain in our *NF1^+/ex42del^* pigs, but a previous study reported that gabapentin or morphine reversed pain behaviors induced by a peripheral neuritis trauma in pigs [7], implying that the two may be linkable. Overall, these data indicates the allodynia and thermal nociception sensitivities detected in the von Frey and laser stimuli are unlikely to be confounded by anxiety or agitation in NF1 mutant pigs.

**Figure 5.**
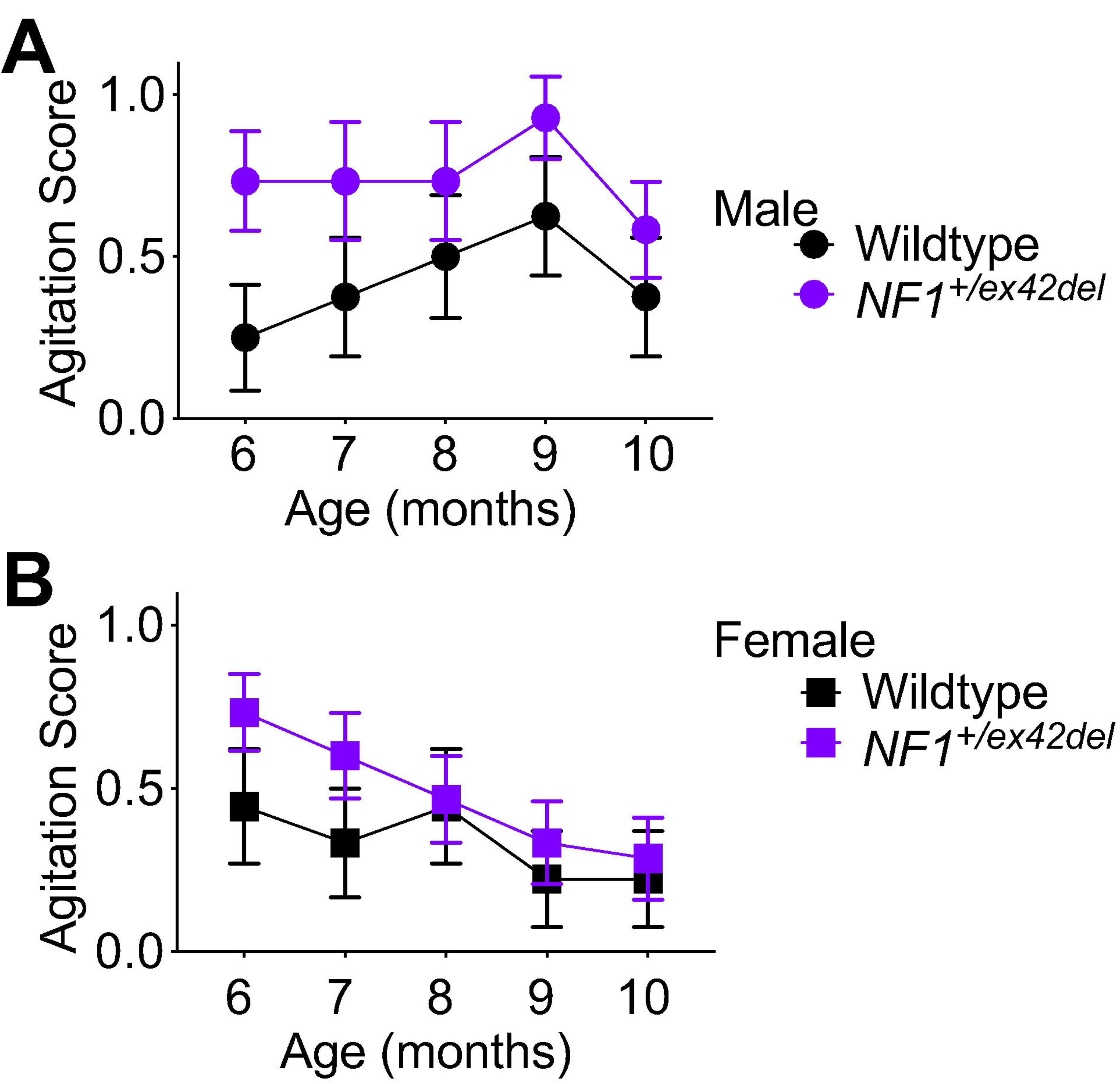
Agitation scores are similar between *NF1^+/ex42del^* and wildtype pigs. Time course of agitation scores for pigs from the indicated genotypes at the indicated age (6 to 10 months). Agitation scores were not significantly different between the genotypes (p>0.05, Mann Whitney for each age).

### 3.4. NF1^+/ex42deI^ pigs are affected in health-related quality of life domains

It has been previously shown that pain is associated with various comorbidities in health-related quality of life domains in NF1 patients [16]. For this reason, we used a FitBark monitoring device to assess sleep quality, and activity related measures (Figure 6) such as percent of day active, resting, and playing, as well as total distance traveled by wildtype and *NF1^+/ex42del^* pigs. While no significant differences were found between wildtype and *NF1^+/ex42del^* pigs in measures of percent of day active (Figure 6A), playing (Figure 6B) or total distance traveled (miles) (Figure 6C), the percent of day resting (Figure 6D) and sleep quality (Figure 6E) were adversely affected in *NF1^+/ex42del^* pigs compared to their wildtype counterparts. Male *NF1^+/ex42del^* pigs with tumors and female *NF1^+/ex42del^* pigs rested longer than wildtype pigs (Figure 6D). Consistent with this data, the sleep quality was reduced in the same male *NF1^+/ex42del^* pigs with tumors and female *NF1^+/ex42del^* pigs compared to wildtype pigs (Figure 6D). These data establish disturbances in health-related quality of life domains in *NF1^+/ex42del^* pigs, consistent with reports of patients with NF1 with mental health and sleep issues and an overall poorer life satisfaction index [16].

**Figure 6.**
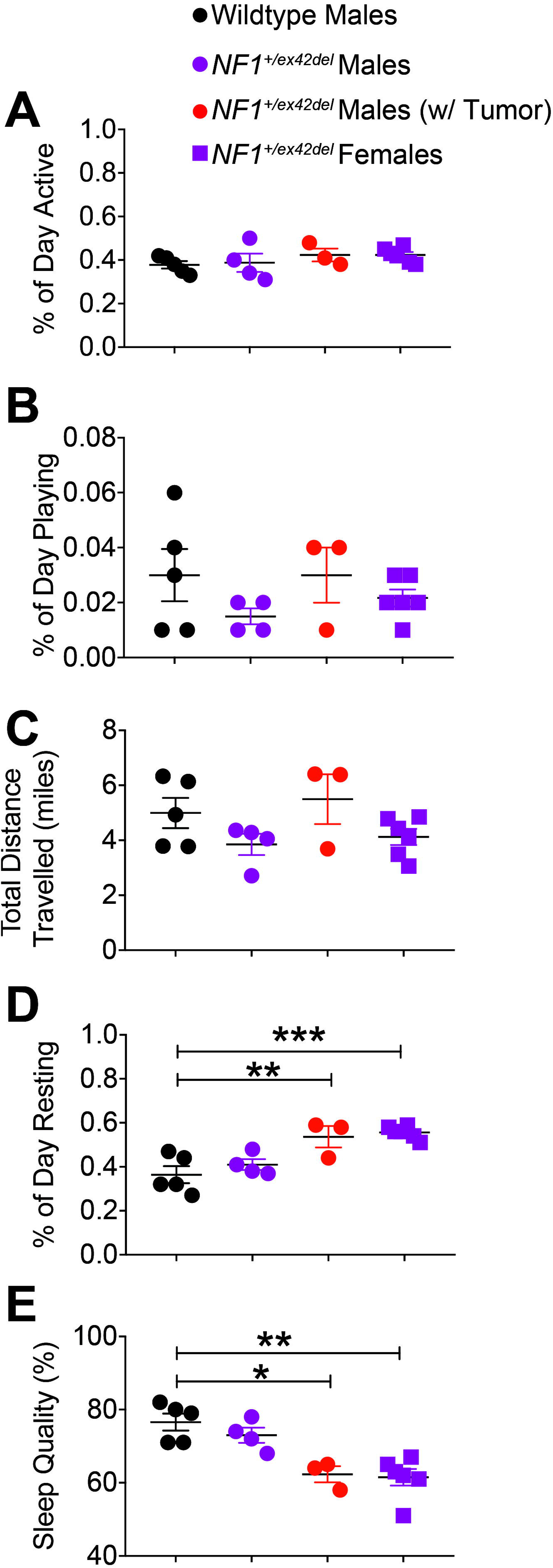
*NF1^+/ex42deI^* pigs exhibit poor sleep and increased time resting. Scatter plots for(**A**) percent of day active, (**B**) percent of day playing, (**C**) total distance travelled, (**D**) percent of day resting, and (**E**) sleep quality for the indicated genotype and gender at 10-12 months of age. Measures of both sleep quality and percent of day resting are adversely affected in female and tumor-burdened male *NF1^+/ex42del^* pigs. Asterisks indicate significance between compared groups (*p<0.05, **p<0.01, ***p<0.001. Oneway ANOVA with Tukey post-hoc).

### 3.5. Increased depolarization of evoked calcium influx in sensory neurons of NF1^+/ex42del^ pigs can be reduced by inhibition of CRMP2 phosphorylation

In NF1, the expression and function of voltage-gated calcium channels is increased [41; 55]. This increase was demonstrated to be reliant on CRMP2 phosphorylation level (S522) in rodent models [41]. Since we found increased CRMP2 phosphorylation at the Cdk5 site (S627) in *NF1^+/ex42del^* pig (Figure 2B), we next asked if the CRMP2 phosphorylation inhibitor (S)-lacosamide ((S)-LCM) [35; 36; 39] could be used to reverse the increased calcium influx as we previously demonstrated in a rat model of NF1 [41]. DRG cultures were generated from either wildtype or *NF1^+/ex42del^* pig DRGs and then imaged for depolarization- evoked calcium influx. We observed that *NF1^+/ex42del^* pig DRGs had an augmented calcium influx compared to wildtype pig DRGs (Figure 7), thus replicating our previous finding in pig [55] and in rodent [41] neurons. Treatment with (S)-LCM decreased depolarization-evoked calcium influx in both wildtype and *NF1^+/ex42del^* pig DRGs (Figure 7). These data demonstrate that pharmacological antagonism of CRMP2 phosphorylation can reduce the activity of voltage-gated calcium channels, setting the stage for translational studies to evaluate this small molecule in future pain behaviors in this NF1 mutant miniswine model.

**Figure 7.**
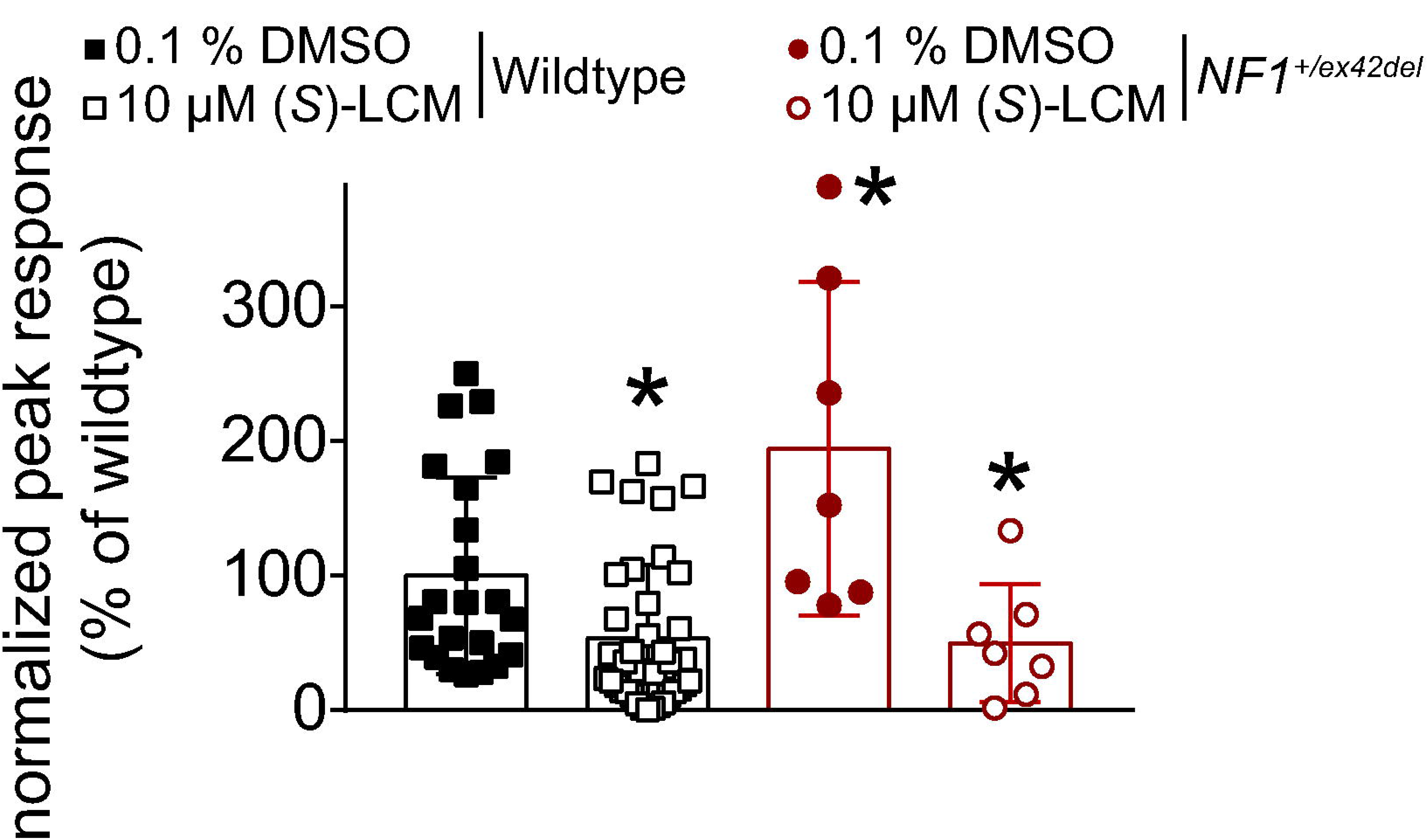
Pharmacological antagonism of CRMP2 phosphorylation by (S)-lacosamide ((S– LCM) reverses the enhanced calcium influx observed in *NF1^+/ex42del^* pig DRG neurons. Bar graph with scatter plot of fura-2 based calcium imaging on primary cultures from *NF1^+/ex42del^* or wildtype pig DRG neurons. After a 1-min baseline acquisition, calcium influx was evoked by a 15 seconds depolarization using 90 mM KCl. Peak response is the difference of peak calcium influx normalized to its own baseline for each individual cell. Peak responses were normalized to the wildtype levels for better comparison. Vehicle (0.1% DMSO) or 10 μM (S)-LCM were applied for 30 min prior to depolarization. Peak calcium response was increased in *NF1^+/ex42del^* pig DRGs compared to wildtype. (S)-LCM treatment inhibited calcium influx in both genotypes. *p<0.05 (Mann Whitney) compared to wildtype (0.1 % DMSO).

## 4. Discussion

Research spanning the last thirty years has shown that the heterogeneity of clinical manifestations of NF1 is not seen with rodent models, limiting their translational impact. We recently described a NF1 mutant miniswine model that exhibited clinical hallmarks of NF1, including café-au-lait macules, neurofibromas, learning and social interaction deficits, as well as remodeling of nociceptive ion channel signaling [55]. Here, we demonstrated: (*i*) increases in voltage-gated calcium channels that are attributable to increased phosphorylation of CRMP2, a known allosteric regulator of these channels [9; 17]; *(ii)* first characterization of pain-related behaviors in a swine model which allowed identification of lower tactile (allodynia) and thermal (hyperalgesia) thresholds in *NF1^+/ex42del^* pigs compared to their wildtype counterparts; *(iii)* disrupted sleep quality and greater rest time in *NF1^+/ex42del^* pigs; and (iv) normalization of increased calcium influx in *NF1^+/ex42del^* pig DRGs by pharmacological antagonism of CRMP2 phosphorylation. Overall, the data supports the use of the *NF1^+/ex42del^* pigs as an improved model for studying NF1 and for translational studies compared to rodents and suggests that interfering with CRMP2 phosphorylation might be a promising therapeutic strategy for NF1-related pain management.

An issue in interpreting signaling changes in mutant *NF1* miniswine is delineating the mechanism of increased calcium signaling and its possible relationship to pain. In our earlier studies with deletion of the *Nf1* gene in rodents, we had identified a signaling pathway that involved increased phosphorylation of CRMP2, a regulator of N-type CaV2.2 channels. CRMP2 bound to CaV2.2 leading to increased Ca^2+^ current density and increased neurotransmitter release in sensory neurons [9]. CRMP2 interacted with the C-terminal domain of neurofibromin [28; 43], such that deletion of neurofibromin (or the C-terminal portion) increased CRMP2 phosphorylation [43], which in turn, increased its association with CaV2.2 [6; 36]. *Nf1* deletion was accompanied by an increase in sensory neuron excitability, and increased depolarization-evoked calcitonin gene-related peptide (CGRP) release in the spinal cord [34]. We determined that these dysregulations were due to the loss of CRMP2 interaction with neurofibromin, thus identifying the CRMP2-neurofibromin interface and CRMP2 phosphorylation as drivers of NF1-related pain [38; 41]. The increase in calcium currents in DRGs from *NF1^+/ex42del^* pigs is consistent with our previous report of increased calcium influx measured with calcium-dye based fluorescence methods [55] as well as with reports in mouse [14; 52] and rat [41] models of NF1.

Notably, our unbiased phosphoproteomics data bring to bear new evidence regarding the allosteric regulation of CRMP2. First, we identified a larger variant of CRMP2, which has also been found in humans [47] but absent in rodents. The porcine peptides belonging to the long form of CRMP2, sometimes described as CRMP2-A [1; 59], were detected here by mass spectrometry. This observation highlights a greater degree of similarity between pigs and humans compared to rodents, supporting the notion that pigs are superior models of the human condition than rodents. Thus, to better model a disease related to CRMP2, one needs to use an animal model expressing human-equivalent splice variants, and pigs express several splice variants for CRMP2 [59]. Thus, our *NF1^+/ex42del^* pig model is a more suitable for studying this CRMP2-related genetic disease.

Second, we found three sites of phosphorylation in CRMP2, including two previously unknown phosphorylated residues in pig CRMP2 – S27 and S627. Because these phosphorylation sites were found from whole DRG lysates, it is unknown whether these are neuron specific events or also present in non-neuronal cells. Among these predicted kinases, increased KIS expression level was reported in NF1-associated plexiform neurofibromas and in MPNSTs [4]. ALKs are known regulators of neurofibromin function [19] and participate in learning and cognitive defect in a mouse model of NF1 [53; 54]. Therefore, CRMP2 phosphorylation by KIS or ALK may contribute to NF1 pathophysiology. In relation to pain behaviors in NF1, we have identified increased CRMP2 phosphorylation by Cdk5 (S522 site in rodents, S627 in pigs from this study) to underlie NF1-related pain [41]. We also found that the CRMP2 phosphorylation inhibitor (S)-LCM, reversed the increased calcium influx in *NF1^+/ex42del^* pig DRGs [35; 36]. This shows that the mechanism of increased voltage gated calcium channel function in NF1 involving increased CRMP2 phosphorylation by Cdk5 is translational from rodent models to our mutant pig model.

Recently, the NF1 research community has lobbied for improved models of the disease to better understand the essential molecular mechanisms underlying the pathogenesis of NF1 and contribute to preclinical studies. Pain in NF1 has been largely overlooked [10; 57]. Between 29-70% of NF1 patients report pain [10; 16], with ~70% of children and adults with NF1 using prescription pain medications [10], and pain being reported as a key symptom of NF1 patients affecting their quality of life [10; 32; 49; 57]. Large animal models of rare diseases are important for understanding disease mechanisms and driving therapeutic discovery. However, behavioral profiling in these large, barn-housed species is challenging and not well-characterized. Typically, pain in pigs has been estimated by responses to nociceptive stimuli, using vocalization, or a composite of behavioral parameters such as lameness, aggression, restlessness, posture, isolation, appearance, sling time, agitation, and posture. Other investigators have analyzed pain indicators in terms of behavior such as inactivity, huddling up, trembling, tail-wagging, scratching, stiffness, sleep spasms, recumbency, coprophagy, aggression, depression, head-pressing, changes in activity, nursing, lying, body movement, muscle-twitching, and withdrawal [18]. In a pig model for chronic pain caused by surgically induced peripheral neuritis trauma, Castel and colleagues [7] used von Frey filaments to assess mechanical sensitivity, a pigeon feather to asses tactile sensitivity, and a composite behaviors scale to assess spontaneous pain behaviors. We found that both female and tumor-burdened male *NF1^+/ex42del^* pigs responded to lower target forces of mechanical stimulation than their wildtype counterparts, suggesting presence of allodynia. We also observed a hyperalgesic phenotype in *NF1^+/ex42del^* pigs, when thermally stimulated, which is consistent with hyperalgesia noted in mice haploinsufficient for *Nf1* [31]. Remarkably, this hyperalgesic behavior was observed only in female pigs, suggesting a sexual dimorphism in pain in NF1. Agitation scores were unchanged between *NF1^+/ex42del^* pigs compared to their wildtype counterparts, indicating that agitation did not play a role in sensitivity to mechanical or thermal stimuli.

As pain and sleep deficits are also observed in NF1 patients [16], it is remarkable that *NF1^+/ex42del^* male (with tumors) and female pigs exhibit depreciated sleep quality in comparison to their wildtype counterparts. Along the same lines, the *NF1^+/ex42del^* male pigs (with tumors) and female pigs spent a greater percent of the day resting than the wildtype pigs housed under identical conditions. As we were able to differentiate these quality-of-life-related measures in the male *NF1^+/ex42del^* pigs on the basis of presence or absence of tumors, and these data show that the affected male *NF1^+/ex42del^* pigs are those also exhibiting tumors, we propose that these pain comorbidities may be related to the presence of tumors in the *NF1^+/ex42del^* pigs. However, a limitation of this work is that the miniswine cohort available to us did not allow us to separate the female *NF1^+/ex42del^* pigs on the basis of presence or absence of tumors because, to date, none of the females have developed tumors. Future studies that include female *NF1^+/ex42del^* pigs with tumors will provide a more complete dataset that may elucidate further gender and tumor-specific phenotypic differences on the basis of pain comorbidities in health-related quality of life domains.

From a clinical perspective, pain is comorbid with symptoms in many health-related quality-of-life domains. Even with regard to the general, non-NF1 population, pain has often been seen in conjunction specifically with depreciated sleep quality. For example, in approaching therapies for chronic spinal pain, Malfliet and colleagues also addressed comorbid insomnia [30]. Song and colleagues highlighted the correlation between migraine and poor sleep quality [46]. In fact, ~48% of the 2695 participants with reported migraines also exhibited poor sleep quality. Along these lines, pain is comorbid with sleep deficits in the NF1 patient population as well. Fjermestad and colleagues reported that in NF1 patients, there is a positive correlation between pain and sleep problems (0.48) [16]. Our miniswine model of NF1 phenocopies this comorbidity of NF1 pain, allowing us to employ a biopsychosocial approach to NF1 pain research that, to our knowledge, is not possible in any rodent model.

In summary, this work characterizes a novel transgenic miniswine model of NF1 pain. We established several different measures for nociceptive dysregulation such as CRMP2 phosphorylation, voltage gated calcium channel regulation, pain related behaviors, and quality of life-related health measures. Besides providing new insights into mechanisms of NF1 nociception, this pig model of NF1 also captures comorbid symptoms of NF1 pain such as sleep deficits that are not seen in rodent models. For these reasons, we propose that the *NF1^+/ex42del^* pig is the most advanced and relevant animal model to study NF1 associated pain.

## Acknowledgements

This work was supported by National Institutes of Health Awards (1R01NS098772, 1R01DA042852, and R01AT009716 to RK); a Neurofibromatosis New Investigator Award from the Department of Defense Congressionally Directed Military Medical Research and Development Program (NF1000099 to RK); and funding from the Synodos for NF1 program at the Children’s Tumor Foundation to DKM, BWD, CSR, JCS, DEQ, and JMW; a research award from the Children’s Tumor Foundation (2015-04-009A) to RK and JMW; a Children’s Tumor Foundation NF1 Synodos award to R.K. A.M. was supported by a Young Investigator’s Award from the Children’s Tumor Foundation. SSB was supported by funds to the University of Arizona’s Undergraduate Biology Research Program. The authors would like to thank Trisha Smit, Brian Dacken, and Justin Van Kalsbeek of Exemplar Genetics, Iowa, USA for their assistance in swine behavior testing and management.

