## Supplementary material for "Sex-dependent differences in pain and sleep in a porcine model of Neurofibromatosis type 1": Table 1

**Table 1. Description of tumors in the *NF1^+/ex42del^* pigs.**

| ID | Sex | DOB | Genotype | Age at Tumor Initiation (mo) | Tumor Description |
| --- | --- | --- | --- | --- | --- |
| A6160 | Boar | 11/18/17 | *NF1^+/ex42del^* | 7 | Small bump on left middle side about 4cm diameter, raised about 0.5cm |
| A6179 | Boar | 11/22/17 | *NF1^+/ex42del^* | 7 | Large bump on left side about 11cm x 7cm – raised 1-2cm |
| A5598 | Boar | 10/18/17 | *NF1^+/ex42del^* | 8 | One large bump in left middle, odd shaped, about 15cm x 7cm – raised about 2cm; One small bump on right shoulder – about 2cm diameter – raised about 0.5cm |
| A5664 | Boar | 10/24/17 | *NF1^+/ex42del^* | 8 | Medium/larger bump on left middle side about 10cm x 5cm – raised about 1cm |
| A6159 | Boar | 11/18/17 | *NF1^+/ex42del^* | 8 | Small bump on middle left side about 2 cm in diameter – raised about 0.5cm |
| A5524 | Boar | 9/19/17 | *NF1^+/ex42del^* | 9 | small bumps - one L middle side, one L shoulder, both about 2cmx2cm, raised 1 cm |
| A5725 | Boar | 11/7/17 | *NF1^+/ex42del^* | 9 | Small bump on right shoulder about 2cm diameter – raised 0.5-1cm |
| A6173 | Boar | 11/20/17 | *NF1^+/ex42del^* | 9 | Bump - back L side (has gotten bigger) |
| A5465 | Boar | 9/11/17 | *NF1^+/ex42del^* | 12 | Bump - middle of spine |
| A5601 | Boar | 10/18/17 | *NF1^+/ex42del^* | 12 | Small bump - back L of spine |
